## Supplemental for "Rhythmic Sampling and Competition of Target and Distractor in a Motion Detection Task"

### Supplementary Materials

In this Supplementary Materials, we considered several issues related to this study: (1) signal processing issues underlying the spectral analysis of various time series data, (2) quantifying distractor processing with MVPA decoding, (3) effect of moving window parameters, and (4) robustness of decoding analysis.

##### *Signal processing issues*

The power of the 4.29 Hz (the target) and 6 Hz (the distractor) signals are both modulated at around 1 Hz. In the Fourier spectrum the sidebands should be visible at around 3.29 Hz and 5.29 Hz (4.29 Hz ± 1 Hz) as well as at around 5 Hz and 7 Hz (6 Hz ± 1 Hz). However, there are no clear peaks visible at these frequencies from Figure 2(B). We examine the reason here.

For clean sinusoidal signals with periodic amplitude modulation, we should observe sidebands. However, biological data is noisy, and the SSVEP from each subject shows significant variability in signal-to-noise ratio (SNR) (see definition below). SNR determines whether we can observe sideband frequencies or not. We demonstrate this point first through simulation and then on our data.

(a) Simulation. Simulated signals were generated by adding a 4.29 Hz component and a 6 Hz component together. These were the same frequency components as in our experiment and also, similar to what we observed in the data, the magnitude of the 6 Hz component was made ½ that of the 4.29 Hz component. We then modulated the amplitude of these components at 1 Hz (the same as that observed in our data). The time course is shown in the left panel of Figure S1(A). In the right panel of Figure S1(A), we showed the Fourier spectrum, where the sidebands for both the 4.29 Hz component (at 3.29 Hz and 5.29 Hz) and the 6 Hz component (at 5 Hz and 7 Hz) are clearly seen. Note that there was no noise in this case.

Next, we added noise to the same signal. The signal-noise-ratio is defined as:

$$SNR_{dB} = 10 \cdot log_{10} \frac{P_{signal}}{P_{noise}}$$

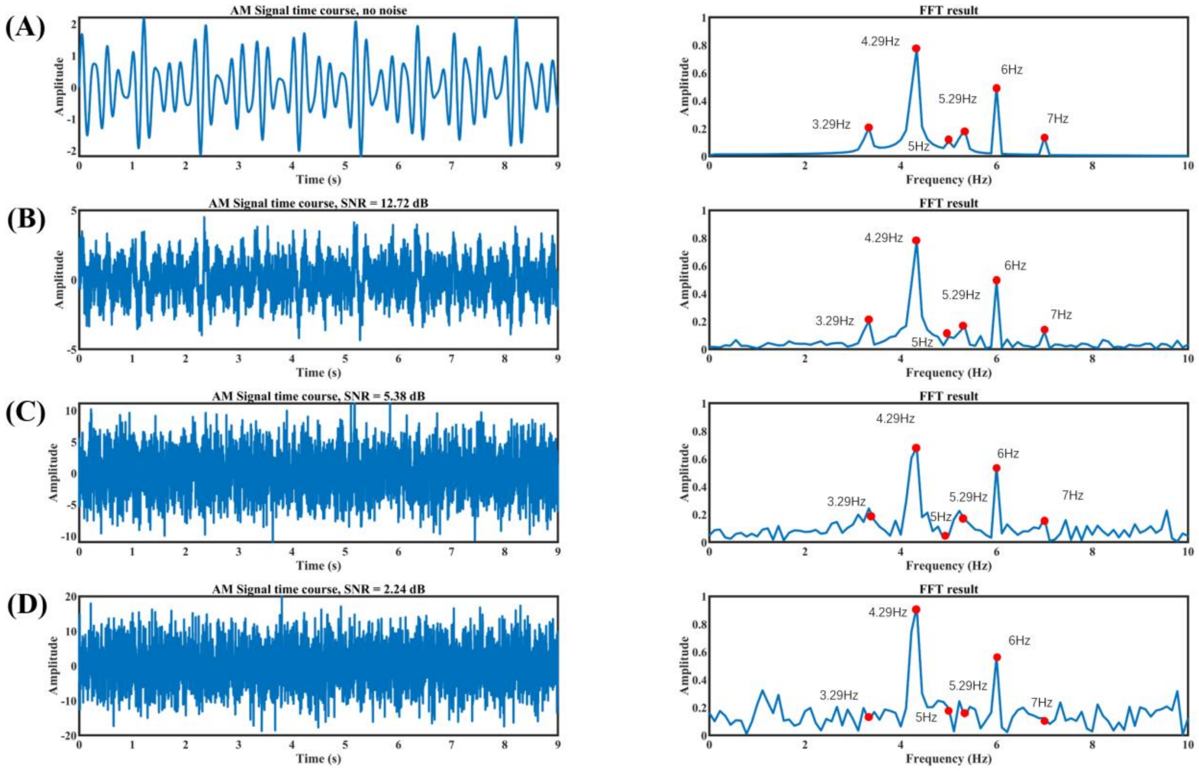


**Figure S1.** Simulation results. (A) The signal containing a 4.29 Hz component and a 6 Hz component where the 6Hz signal's magnitude is about half that of the 4.29Hz signal. The amplitude is modulated at 1 Hz. No noise is added. (B) Low level of noise is added to the signal in Figure S1(A) where the SNR = 12.72 dB. Sidebands are still seen. (C) Middle level of noise is added to the signal in Figure S1(A) where the SNR = 5.38 dB. Sidebands become difficult to see. (D) High level of noise is added to the signal in Figure S1(A) where the SNR = 2.24 dB, sidebands become more indistinguishable from the noise floor. Red dots indicate the location of the main frequency components and the locations where the sidebands should appear.

where the signal power is defined to be the average Fourier power within 3.8 to 4.8Hz and 5.5 to 6.5Hz and the noise power is the average Fourier power within 0 to 3Hz and 7 to 10Hz. Figure S1(B), Figure S1(C), and Figure S1(D) show the results after adding progressively more noise to the simulated signal. When the noise level is low, e.g., SNR=12.72 dB (Figure S1(B)), the sidebands are still clearly visible, as shown in the right panel of Figure S1(B), although they are not as prominent as in the right panel of Figure S1(A). When more noise is added, as shown in Figure S1(C) where SNR=5.38 dB, which is similar to what we see in a typical high SNR subject

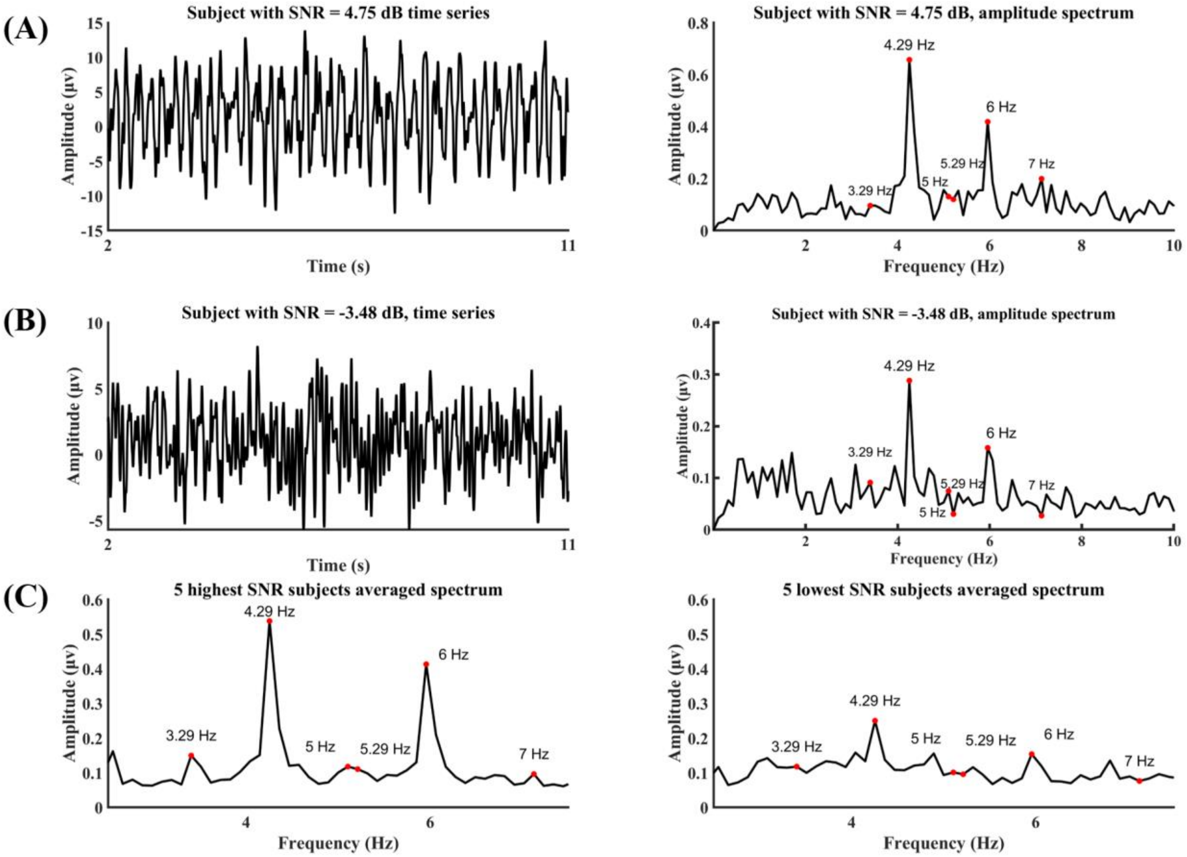


**Figure S2.** Experimental data. (A) The time course of the SSVEP and its Fourier spectrum from a subject with high SNR. The sidebands can be observed. (B) The time course and its Fourier spectrum from a subject with low SNR. The sidebands are indistinguishable from the noise floor. (C) The averaged Fourier spectrum from 5 highest SNR subjects and 5 lowest SNR subjects. Again, for subjects with high SNR, the sidebands are identifiable, whereas for subjects with low SNR, the sidebands are not identifiable.

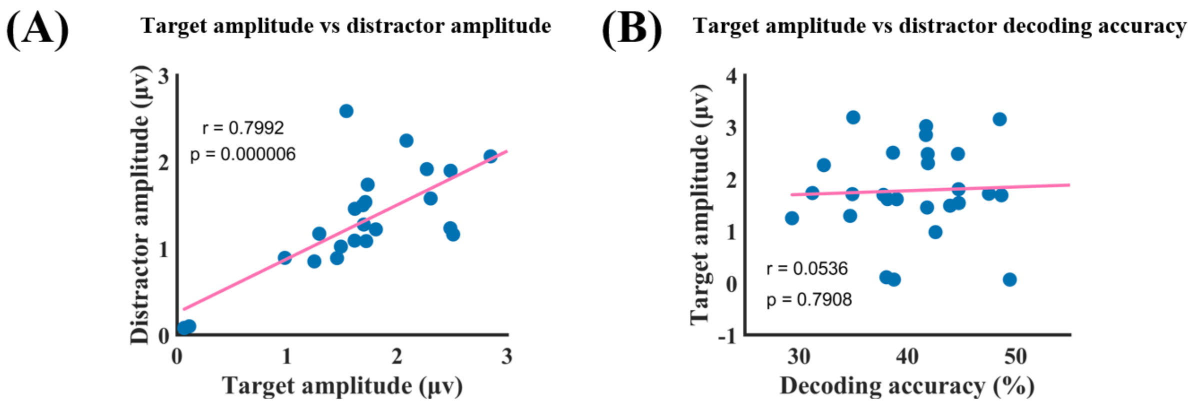


**Figure S3.** SSVEP amplitude analysis at the whole trial level. (A) Target amplitude vs distractor amplitude, where the correlation is r = 0.7992 (p = 0.000006), suggesting the 6 Hz signal amplitude is strongly influenced by the 4.29 Hz signal amplitude. (B) Target amplitude vs distractor decoding accuracy, where the correlation is r = 0.0536 (p = 0.7908), suggesting that the decoding accuracy as an index of distractor processing is not influenced by the 4.29 Hz target amplitude.

*(b) Moving window analysis.* Similar results can be observed at the moving window level. The target amplitude time series and the distractor amplitude time series were extracted from each subject and the relation between the two time series assessed. The relative phase is also correlated

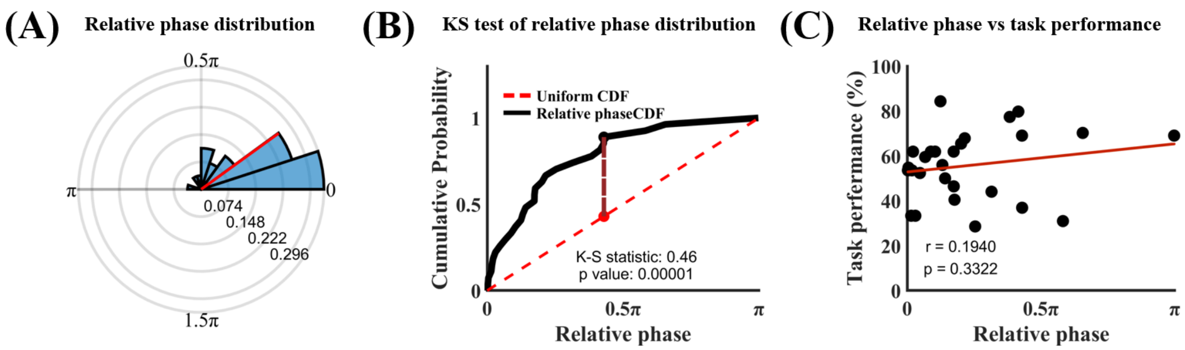


**Figure S4.** Moving window analysis. (A) The relative phase between the target amplitude time series and the distractor amplitude time series. (B) Kolmogorov-Smirnov test showed that the relative phase distribution is significantly different from the uniform distribution. (C) Relative phase vs task performance. r=0.1940 (p=0.3322) means that there is no significant correlation between amplitude relative phase and task performance.

with behavior. Figure S4(A) shows that the relative phase is narrowly distributed around 0.23 ± 0.05π, which is confirmed by the K-S static of 0.46 in Figure S4(B), demonstrating that the relative phase distribution is significantly different from the uniform distribution at p=0.00001. This means that the power leakage from the target time series impacts the distractor time series, making it nearly phase locked to the target time series. It is not surprising that the relative phase between the two amplitude time series does not predict task performance (Figure S4(C)).

***Effect of moving window parameters***

We redid the moving window analysis using a different set of windowing parameters, e.g., a 0.1s sliding window length with a 0.05s step size. Figure S5 demonstrates that the strength of both target and distractor processing fluctuates around ~1 Hz, both at the individual and group levels. Additionally, Figures S6(A) and S6(B) show that the relative phase between target and

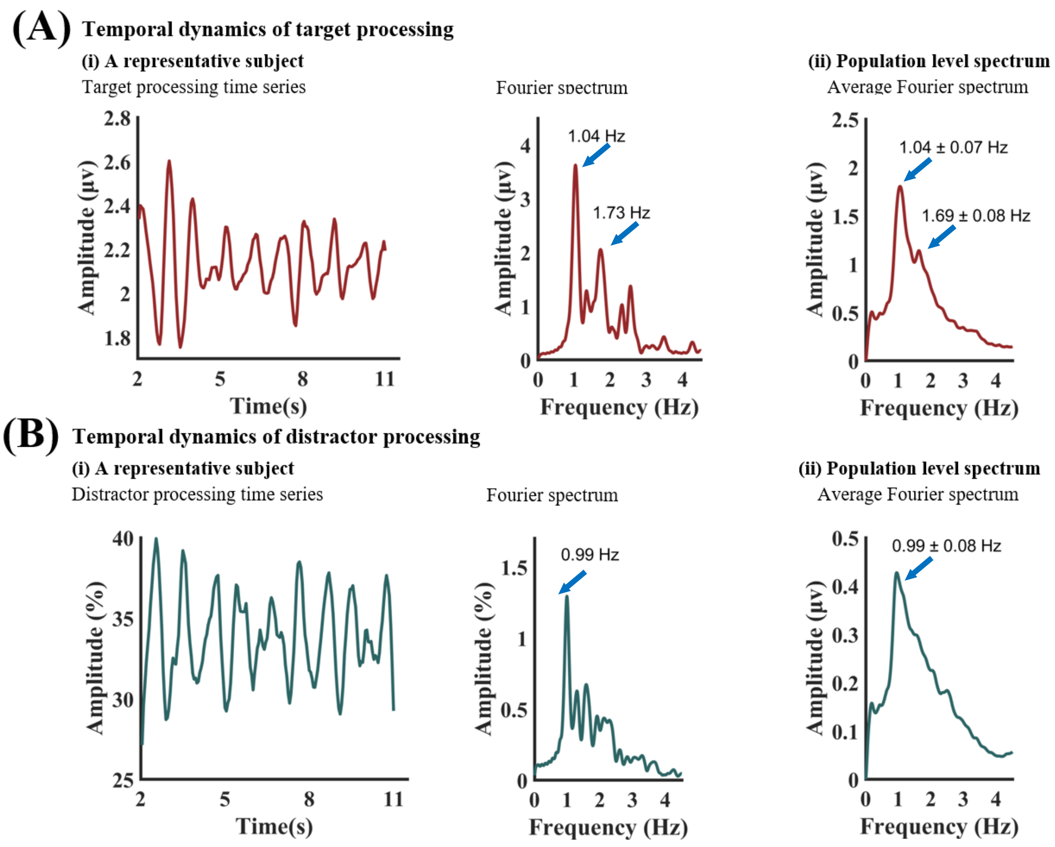


**Figure S5.** Temporal dynamics of target and distractor processing with 0.1s window length and 0.05s step size. (A) (i): Target processing time series from the moving window approach for a representative subject (left) and its Fourier spectrum (right). (A) (ii): The average Fourier spectrum across 27 subjects. (B) (i): Distractor processing time series from the moving window approach for a representative subject (left) and its Fourier spectrum (right). (B) (ii): The average Fourier spectrum across 27 subjects.

distractor processing time series exhibits a uniform distribution across subjects. For the relation between relative phase and behavior, Figure S6(C) illustrates two representative cases: a high-performing subject with 84.34% task accuracy exhibited a relative phase of 0.9483π, while a low-performing subject with 30.95% accuracy showed a phase of 0.29π. At the group level, a significant positive correlation between relative phase and task performance was found (r = 0.6343, p = 0.0004), as shown in Figure S6(D). All these results, aligning closely with our original findings, suggest that the conclusions are not dependent on windowing parameters.

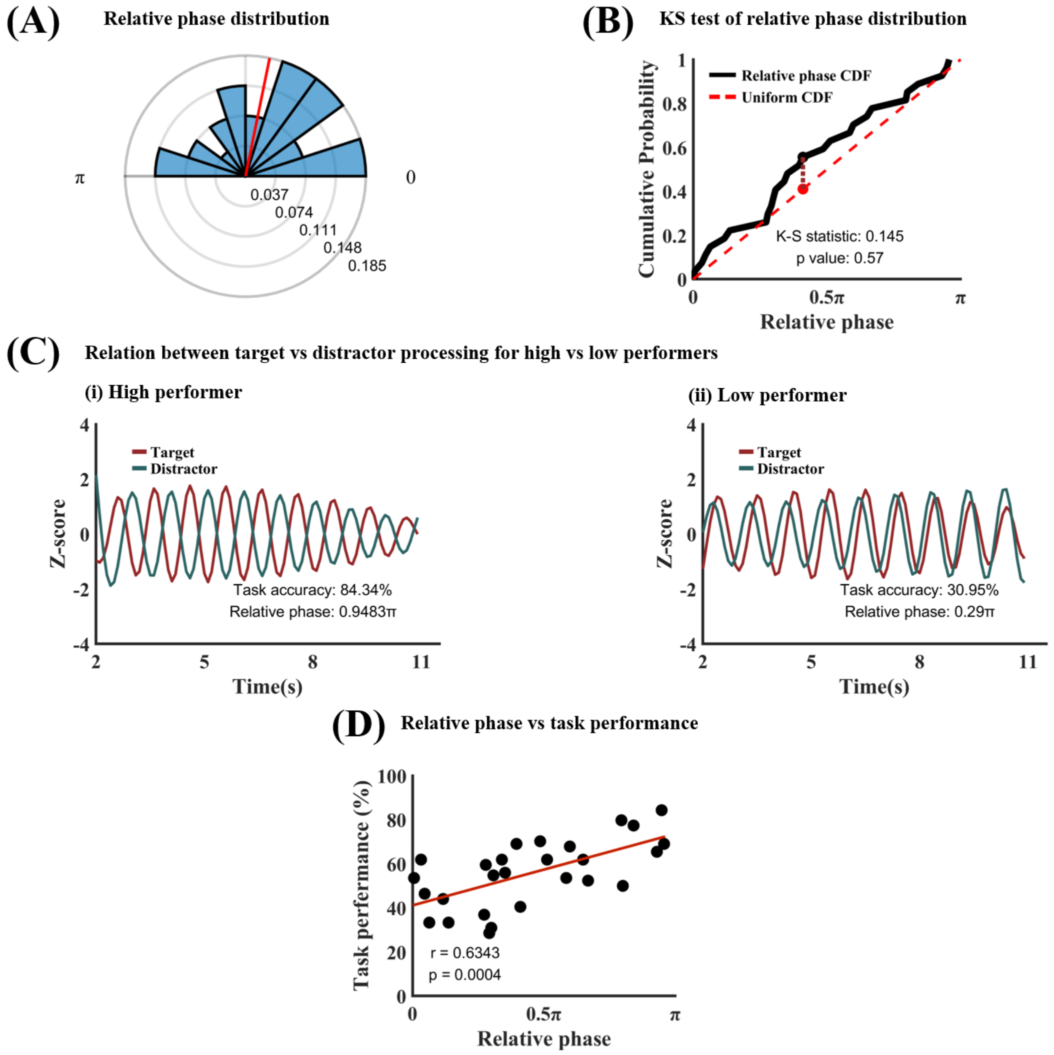


**Figure S6.** Target-distractor competition analysis with 0.1s window length and 0.05s step size. (A) Phase polar histogram for the relative phase between target process time series and distractor processing time series (1 Hz). The average relative phase is 0.44π. (B) Kolmogorov-Smirnov test showed that the relative phase distribution is not different from uniform distribution. (C) Temporal relation between target processing and distractor processing for (i) a high performer (accuracy=83.84%; relative phase=0.9483π) and (ii) a low performer (accuracy=30.95%; relative phase=0.29π). (D) Task performance vs 1 Hz relative phase. The significant positive correlation (r=0.6343, p=0.0004) means that the more separated the target and distractor sampling within the 1 Hz oscillation cycle the better the behavioral performance. CDF: Cumulative distribution function.

To further validate our findings, we also employed the Hilbert transform to extract amplitude envelopes of the target and distractor signals on a time-point-by-time-point basis, providing a window-free estimate of signal strength (Figures S7 and S8). The results remain consistent with both the original findings and the new sliding window analyses (above). Figure S7 reveals ~1 Hz fluctuations in target and distractor processing at both individual and group levels. Figures S8(A) and S8(B) confirm a uniform distribution of the relative phase. As shown in Figure S8(C), the relative phase was 0.9567π for a high-performing subject (84.34% accuracy) and 0.2247π for a low-performing subject (28.57% accuracy). At the group level, a significant positive

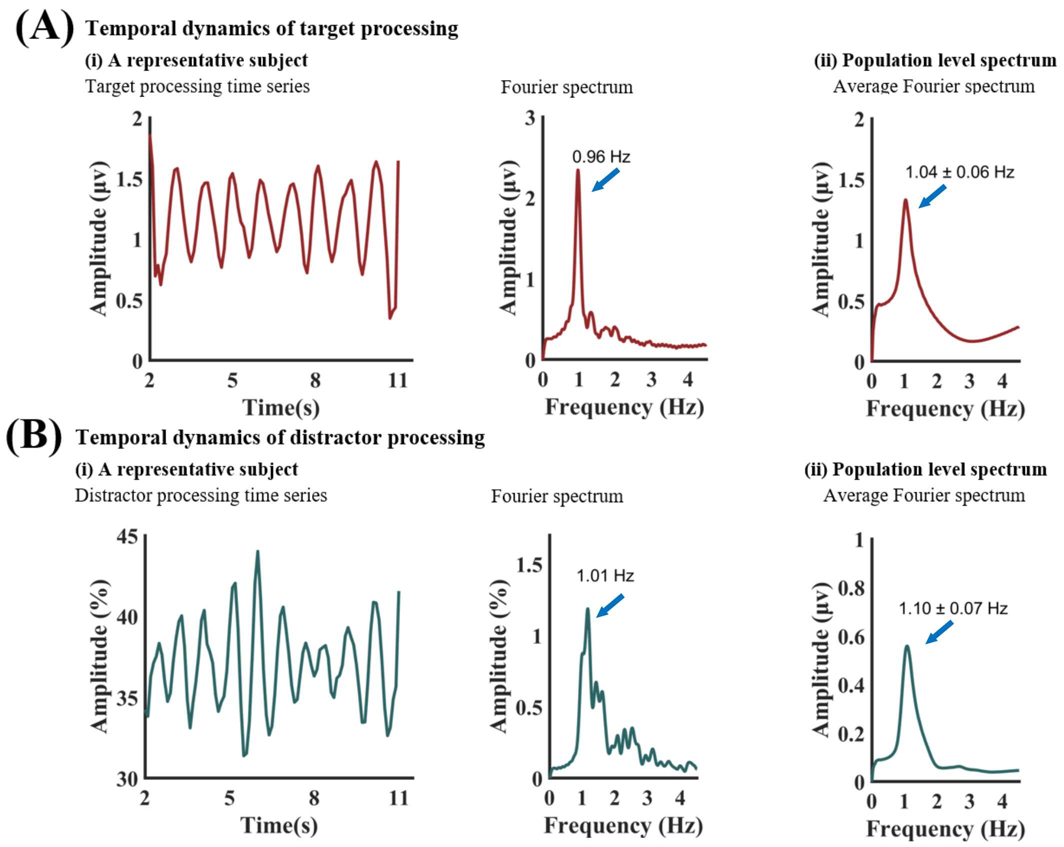


**Figure S7.** Temporal dynamics of target and distractor processing with Hilbert transformed target and distractor processing time series. (A) (i): Target processing time series from for a representative subject (left) and its Fourier spectrum (right). (A) (ii): The average spectrum across 27 subjects. (B) (i): Distractor processing time series for a representative subject (left) and its Fourier spectrum (right). (B) (ii): The average spectrum across 27 subjects.

correlation was again observed between relative phase and task performance (r = 0.4020, p = 0.0376), as shown in Figure S8(D).

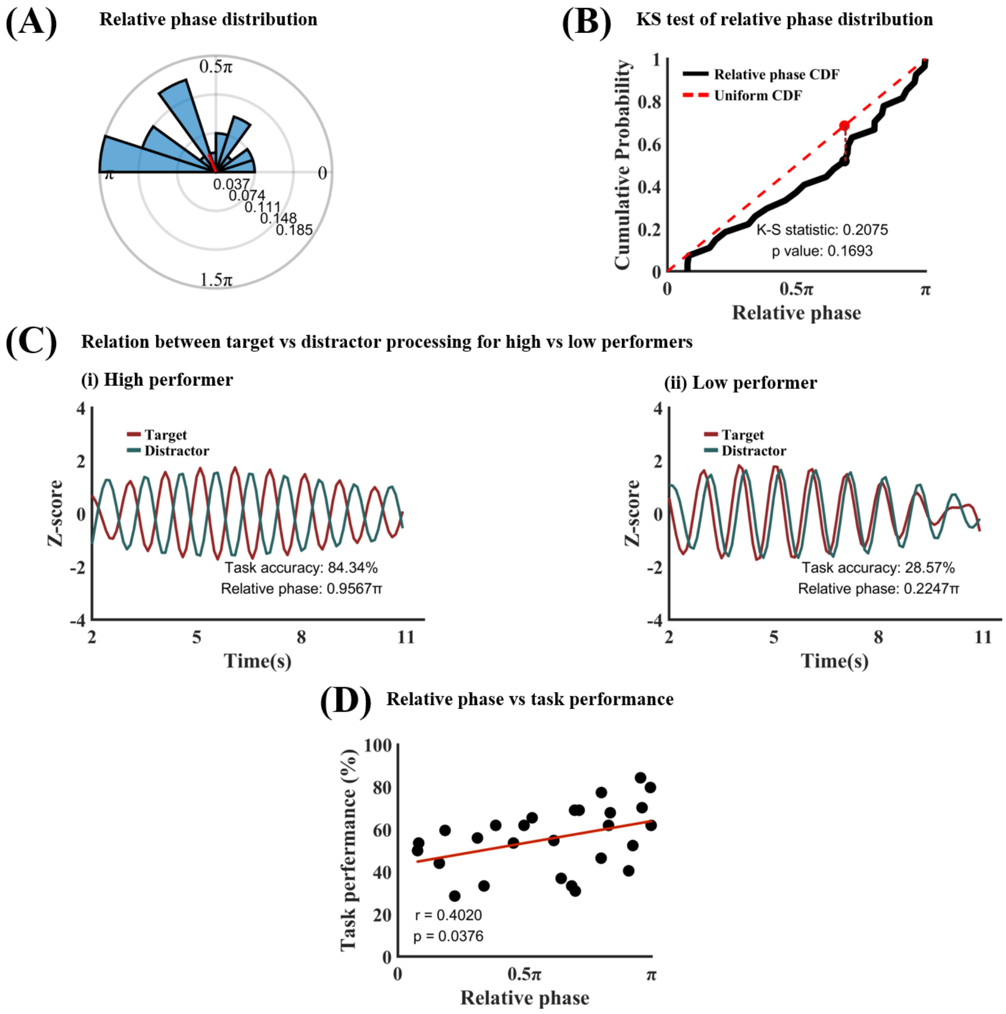


**Figure S8.** Target-distractor competition analysis with Hilbert transformed target and distractor processing time series. (A) Phase polar histogram for the relative phase between target process time series and distractor processing time series (1 Hz). The average relative phase is 0.63π. (B) Kolmogorov-Smirnov test showed that the relative phase distribution is not different from uniform distribution. (C) Temporal relation between target processing and distractor processing for (i) a high performer (accuracy=83.84%; relative phase=0.9567π) and (ii) a low performer (accuracy=28.57%; relative phase=0.2247π). (D) Task performance vs 1 Hz relative phase. The significant positive correlation (r=0.4020, p=0.0376) means that the more separated the target and distractor sampling within the 1 Hz oscillation cycle the better the behavioral performance. CDF: Cumulative distribution function.

##### *Robustness of decoding analysis*

To test robustness of the decoding analysis, we implemented a random permutation procedure in which trial labels were randomly shuffled to construct a null-hypothesis distribution of decoding accuracy. We then compared the decoding accuracy from the actual data to this distribution. Figure S9 shows the results based on 1,000 permutations. For each of the three pairwise classifications—pleasant vs. neutral, unpleasant vs. neutral, and pleasant vs. unpleasant—as well as the three-way classification, the actual decoding accuracies fall far outside the null-hypothesis distribution ($p < 0.001$), and the effect sizes are extremely large. These findings indicate that the observed decoding accuracies are statistically significant and robust in terms of both statistical inference and effect size.

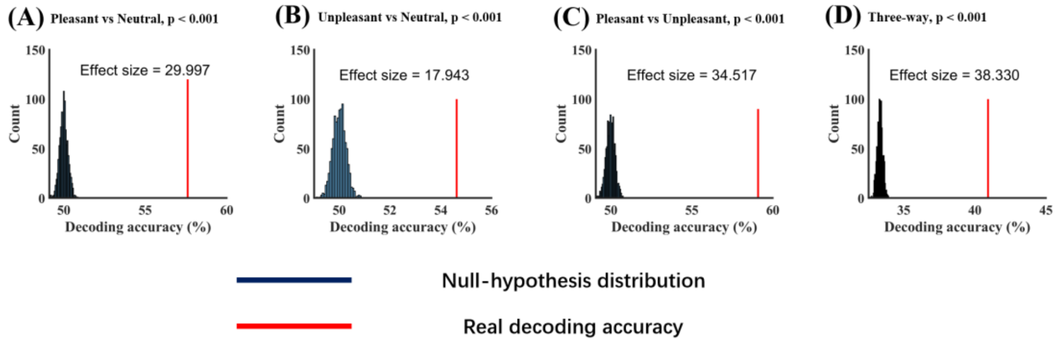


**Figure S9.** Comparison of actual decoding accuracy against the distribution of random permutation decoding accuracy. Random permutation decoding accuracy from (A) Pleasant vs Neutral, (B) Unpleasant vs Neutral, (C) Pleasant vs Unpleasant, and (D) Three-way. In all four conditions, the actual decoding accuracy is significantly above chance level at $p<0.001$.
